# Emergence and spread of SARS-CoV-2 variants from farmed mink to humans and back during the epidemic in Denmark, June-November 2020

**DOI:** 10.1101/2024.02.13.580053

**Authors:** Thomas Bruun Rasmussen, Amanda Gammelby Qvesel, Anders Gorm Pedersen, Ann Sofie Olesen, Jannik Fonager, Morten Rasmussen, Raphael Niklaus Sieber, Marc Stegger, Francisco Fernando Calvo-Artavia, Marlies Jilles Francine Goedknegt, Esben Rahbek Thuesen, Louise Lohse, Sten Mortensen, Anders Fomsgaard, Anette Boklund, Anette Bøtner, Graham J. Belsham

## Abstract

The severe acute respiratory syndrome coronavirus-2 (SARS-CoV-2) not only caused the COVID-19 pandemic but also had a major impact on farmed mink production in several European countries. In Denmark, the entire population of farmed mink (over 15 million animals) was culled in late 2020. During the period of June to November 2020, mink on 290 farms (out of about 1100 in the country) were shown to be infected with SARS-CoV-2. Genome sequencing identified changes in the virus within the mink and it is estimated that about 4000 people in Denmark became infected with these mink virus variants. However, the routes of transmission of the virus to, and from, the mink have been unclear. Phylogenetic analysis revealed the generation of multiple clusters of the virus within the mink. Detailed analysis of changes in the virus during replication in mink and, in parallel, in the human population in Denmark, during the same time period, has been performed here. The majority of cases in mink involved variants with the Y435F substitution and the H69/V70 deletion within the Spike (S) protein; these changes emerged early in the outbreak. However, further introductions of the virus, by variants lacking these changes, from the human population into mink also occurred. Based on phylogenetic analysis of viral genome data, we estimate, using a conservative approach, that about 17 separate examples of mink to human transmission occurred in Denmark but up to 59 such events (90% credible interval: (39-77)) were identified using parsimony to count cross-species jumps on transmission trees inferred using Bayesian methods. Using the latter approach, 136 jumps (90% credible interval: (117-164)) from humans to mink were found, which may underlie the farm-to-farm spread. Thus, transmission of SARS-CoV-2 from humans to mink, mink to mink, from mink to humans and between humans were all observed. (298 words)

**Author summary:** In addition to causing a pandemic in the human population, SARS-CoV-2 also infected farmed mink. In Denmark, after the first identification of infection in mink during June 2020, a decision was made in November 2020 to cull all the farmed mink. Within this outbreak, mink on 290 farms (out of about 1100 in the country) were found to have been infected. We showed, by analysis of the viruses from the mink, that the viruses on the farms were mainly of three different, but closely related, types (termed Clusters 2, 3 and 4) that shared certain distinctive features. Thus, we found that many outbreaks in mink resulted from transmission of the virus between mink farms. However, we identified that new introductions of other virus variants, presumably from infected humans, also occurred. Furthermore, we showed that spread of the virus from infected mink to humans also happened on multiple occasions. Thus, transmission of these viruses from humans to mink, mink to mink, from mink to humans and between humans were all observed. (172 words)

## Introduction

The severe acute respiratory syndrome coronavirus-2 (SARS-CoV-2) has caused the COVID-19 pandemic [1], with over 675 million cases reported globally and it has contributed to the deaths of at least 6.8 million people [2]. The coronavirus (RaTG13), which has been found to be the most closely related to SARS-CoV-2, was detected in horseshoe bats (*Rhinolophus affinis*) in China [3], with about 1200 nucleotide (nt) differences between their full-length RNA genomes of about 30,000 nt (ca. 96% identity). It is not known how the virus moved from these bats to humans or if there was an intermediate host [4, 5], as with civet cats for the SARS-CoV [6]. In addition to the effect of the continuing pandemic in humans, the same virus has also had a drastic impact on farmed mink production worldwide. Outbreaks of disease on mink farms, caused by infection with SARS-CoV-2, were initially identified, during April 2020, in the Netherlands (NL) [7]. These were followed closely (from June 2020) by outbreaks in Denmark (DK) [8], a country with one of the highest levels (about 40%) of global mink production, involving at that time over 1100 farms and a population of about 17 million mink [9]. Spread of SARS-CoV-2 into mink was also observed in a variety of other countries, including Canada, France, Greece, Italy, Lithuania, Spain, Sweden and the USA [10].

In total, SARS-CoV-2 infections were detected on 290 mink farm premises in DK (ca. 25% of the total) and this contributed to the Danish government’s decision in early November 2020 to stop all mink production within DK [11]. The entire mink population was culled [9] and mink production halted until the end of 2022. The production of mink in the NL was also stopped in 2020, bringing forward an earlier planned end to this industry [12].

During the course of the outbreaks in mink in DK, a large number of different virus variants were observed. However, most of the viruses from mink that were analyzed had a specific mutation (A22920T) within the gene encoding the Spike (S) protein, resulting in the conservative amino acid substitution Y453F (tyrosine to phenylalanine), which occurred on the first mink farm found to have infected animals in DK [8]. This mutation was one of the defining changes that lead to the emergence of the virus pangolin lineage termed B.1.1.298 within the European Clade 20B. This same change was seen on one mink farm in the NL, early in the outbreak there, but also later in other mink farms [7, 13]. However, these variants, belonged to two different clades, 19A and 20A, and did not predominate in the NL. The residue Y453 lies within the receptor binding domain (RBD) of the S protein that is known to interact with the cellular receptor, angiotensin-converting enzyme 2 (ACE2), which is used by the virus [14]. It has been reported that the Y453F substitution enhances binding of the virus to the mink ACE2 protein without compromising interaction with the human ACE2 protein [15].

A second, early, change in the viruses circulating in the mink population was the deletion of six contiguous nucleotides in the S gene coding sequence, which resulted in the loss of two amino acid residues, H69 and V70 (termed H69/V70del), from the S protein [16]. This change was first detected (in August 2020) on the 4^th^ farm with infected mink in DK along with additional sequence changes, in other parts of the virus genome (including nucleotide changes leading to the amino acid substitutions P3395S in ORF1a and S2430I in ORF1b).

After the appearance of the Y453F and H69/V70del variants in mink, viruses with these changes were also found in the human population in the same region of DK, namely Northern Denmark [8, 11]. In total, the mink variants of SARS-CoV-2 were detected in over 1,100 people in DK out of 53,933 sequenced samples during the period from June 2020 to January 2021 [17] and this incidence was used to estimate that about 4000 humans in DK became infected with mink-derived viruses [11]. In Northern Denmark, where most SARS-CoV-2 outbreaks in mink occurred, amongst the people connected to mink farms, about 30% tested positive for SARS-CoV-2 in the period from June to November 2020 and approximately 27% of the SARS-CoV-2 samples from humans in this community were mink-associated [11].

During August and September 2020, mink on substantially more farms tested positive for SARS-CoV-2 [9]. This was coincident with extensive community spread of the virus [11] and further sequence changes generating multiple discrete clusters of viruses (termed Clusters 2, 3, 4 and 5) within the mink phylogeny (Figure 1). There was particular concern about a Cluster 5 isolate (named hCoV-19/Denmark/DCGC-3024/2020, GISAID EPI_ISL_616802), which had a number of amino acid sequence changes in the S protein (Y453F, I692V and M1229I as well as the H69/V70del). Preliminary testing of this virus isolate suggested a possible decrease in neutralization of this virus variant by human antibodies [18]. However, further analysis [19] showed that the impact of these changes on the ability of this virus to be neutralized by antibodies from convalescent humans was generally rather limited. Similarly, it has been found that there was very little loss of neutralization of pseudoviruses carrying a Cluster 5-like S protein, compared to wild-type, by sera from people twice vaccinated with Pfizer or Moderna mRNA vaccines [20].

**Figure 1.**
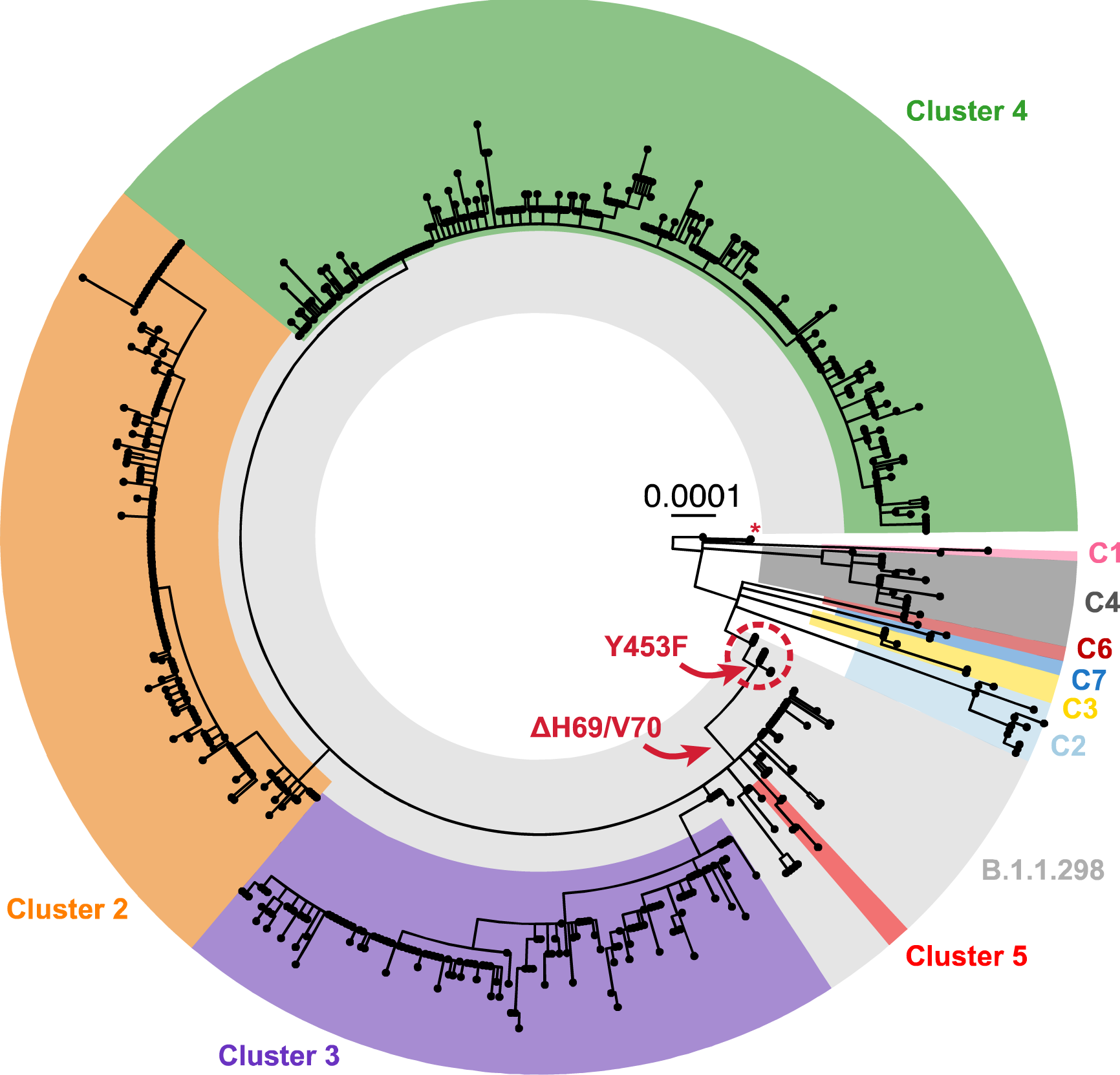
Phylogeny of the 698 SARS-CoV-2 whole-genome sequences from Danish mink. The majority of viruses found on infected farms, including those from the initial cases (farms 1-3, indicated within a red dashed circle) and viruses in Clusters 2-5, belong to pangolin lineage B.1.1.298 and are highlighted in light grey. Clusters 2-5 and viruses subsequently found as further spillovers from humans (C1-C4 and C6-C7) are highlighted in different colors. A singleton sequence belonging to C8 is indicated by a red asterisk. The occurrence of key sequence changes that were present in most mink virus sequences are indicated with red arrows. The scale bar indicates number of substitutions per variable site. The phylogeny was rooted with the basal reference sequence (NC_045512.1, known as the Wuhan-Hu-1 virus) as the outgroup.

On mink farms within Denmark, although many mink became infected with SARS-CoV-2, the incidence of disease was low, with few clinical signs and only low, albeit elevated, levels of mortality [9]. This meant that often the infection was detected at a late stage after infection when most, or frequently all, animals had already seroconverted at first testing. Despite significant effort [9], no potential routes of virus transmission between mink farms, e.g. by wildlife or birds, were identified except for direct contact with infected humans. On one farm, the presence of virus was detected when the prevalence was low (only in 13% of tested mink) and with only 3% seroprevalence in the sampled animals but the virus prevalence had increased to about 90% only 4 days later [8] and the seroprevalence had reached 97% after a further eight days. This indicates a very rapid spread of the virus among the mink that were living in close proximity. Thus the spread of the virus in mink was dependent on rapid transmission within farms in conjunction with undefined routes of transmission between farms.

In the current study, the genomic sequences of viruses from nearly all known infected mink farm premises in DK have been analyzed together with the sequences of the viruses circulating in the human population in DK during the same time period in order to determine the pathways of virus transmission within and between these two host species. The results shed light on the spread and evolution of the virus within mink and identify many occasions when the virus was transmitted from humans to mink, as well as *vice-versa*.

## Results

### Appearance of multiple clusters of SARS-CoV-2 in mink

After the initial cases (starting in June 2020) of SARS-CoV-2 infection on four mink farms in DK [8, 16], there was further spread of the virus to other farms (Figure 1, Table 1). Outbreaks initially occurred within Northern Denmark but spread into Central and Southern Denmark (Figure 2). The virus variants found in mink in DK, during August and September 2020, all belonged to the same pangolin lineage, B.1.1.298, as for the initial cases, and were most likely descendants from the virus identified in the mink population in June. They all had the Y453F substitution in the S protein that was first observed on farm 1 [8]. It should be noted that from farm 1 onwards, each farm with infected mink was numbered consecutively following detection of SARS-CoV-2 on the farm. The SARS-CoV-2 in DK at that time, in both humans and mink, all had the A23403G change (encoding the substitution D614G within the S protein) compared to the Wuhan strain and this change is not considered further.

**Figure 2.**
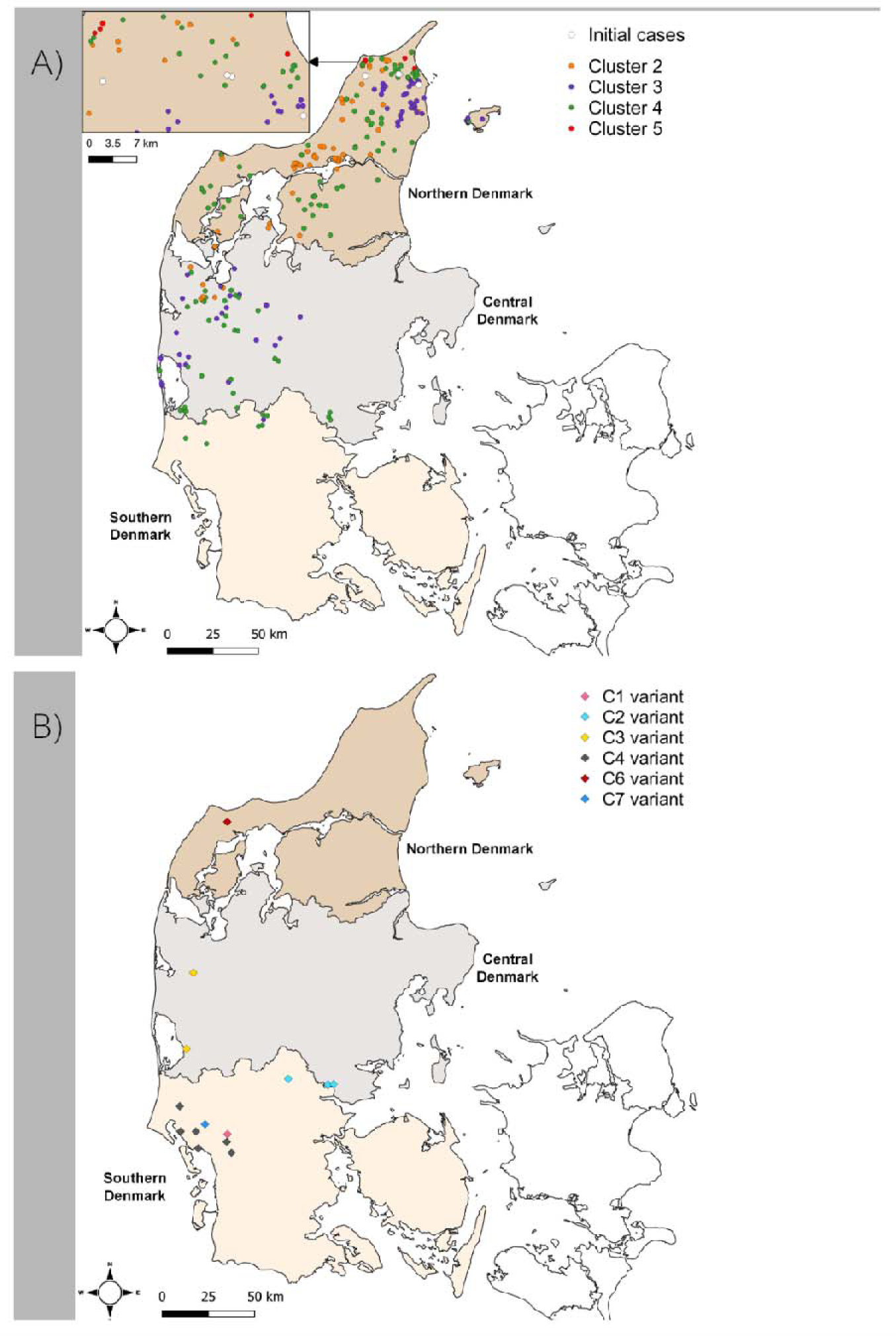
Location of different SARS-CoV-2 variants in mink during the epidemic in Denmark, June-November 2020. Panel A. The location of the initial cases of SARS-CoV-2 infection in Northern Denmark are indicated. Subsequently, further cases occurred and the virus diverged, within lineage B.1.1.298, into Clusters 2, 3, 4 and 5 (as shown in Figure 1). Panel B. Later in the epidemic, new introductions of viruses from different lineages occurred and these are named as C1-C7 (see Table 1). The maps were made using the free and open source geographic information system Q-GIS with the shapefile of the administrative units of Denmark uploaded from DIVA-GIS (open source) onto which the data was plotted, see: https://www.qgis.org/en/site/getinvolved/governance/trademark/index.html

**Table 1.** Summary of the SARS-CoV-2 sequences from infected mink from farms in DK. The features of the different Clusters (as identified in Figure 1) are shown. The later introductions into mink from infected humans, designated C1-C8 are also indicated; these viruses lack the Y453F change.

| Cluster | Pangolin lineage | Clade | Spike protein signature<sup>1</sup> | Spike protein deletion* | No. of mink sequences | No. of farms | First sequence (date) | Last sequence (date) | Location |
| --- | --- | --- | --- | --- | --- | --- | --- | --- | --- |
| **Initial cases** | B.1.1.298 | 20B | (Y435F) | (H69/V70) | 43 | 4 | 14-06 | 06-11 | Northern Denmark |
| **2** | B.1.1.298 | 20B | Y453F | H69/V70 | 174 | 76 | 09-09 | 12-11 | Northern/Central Denmark |
| **3** | B.1.1.298 | 20B | Y453F | H69/V70 | 142 | 66 | 14-09 | 15-11 | Northern/Central Denmark |
| **4** | B.1.1.298 | 20B | Y453F | H69/V70 | 272 | 121 | 10-09 | 03-12 | Northern/Central Denmark |
| **5** | B.1.1.298 | 20B | Y453F | H69/V70 | 5 | 5 | 31-08 | 15-09 | Northern Denmark |
| **C1** | B.1.258.9 | 20A | N439K;<br>G1223S | H69/V70 | 2 | 1 | 03-11 | - | Southern Denmark |
| **C2** | B.1.1.219 | 20B | F157L;<br>(A845S) |  | 13 | 4 | 16-10 | 13-11 | Central Denmark |
| **C3** | B.1.1.170 | 20B | (G1167S) |  | 6 | 3 | 23-10 | 29-10 | Central Denmark |
| **C4** | B.1.536 | 20A | (N501T) |  | 19 | 7 | 16-10 | 18-11 | Southern Denmark |
| **C6** | B.1.1.294 | 20B |  |  | 3 | 1 | 23-10 | - | Northern Denmark |
| **C7** | B.1.1.159 | 20B |  |  | 3 | 1 | 12-11 |  | Southern Denmark |
| **C8** | B.1.177 | 20E | A222V |  | 1 | 1 | 02-11 | - | Northern Denmark |

<sup>1</sup>: Changes shown in parenthesis were only found in some of the mink sequences in the cluster

Additional mutations emerged within the infected mink. Whole-genome-based phylogenetic analysis, using the maximum-likelihood method, performed on 698 sequences from infected mink (from nearly all the affected farms in DK), showed a segregation of the viruses from the initial cases into four major clusters (termed Clusters 2, 3, 4 and 5) indicating multiple independent transmission pathways from farm to farm (Figure 1 and Supplementary Figure 1). As indicated above, it can be envisaged that within a single farm there will be rapid spread of the virus that was introduced into the farm between the mink but onward spread to another farm probably involved a different mechanism of transmission.

A circular representation of the phylogenetic tree clearly shows the general dominance of Clusters 2, 3 and 4 within this epidemic (Figure 1), but sequencing was only performed on a small subset of the infected mink, thus the precise proportions of mink infected with each variant is not known. A rectangular version of the phylogenetic tree based on the same set of virus sequences, but including sequence IDs and farm numbers, is shown in Supplementary Figure S1.

Viruses present on farms 1-4 [16], represent parental sequences to Clusters 2, 3, 4 and 5 (Supplementary Figure S1). In total, 270 of the 290 farms (i.e. 93%) that were tested positive for SARS-CoV-2 by the end of November had mink infected with variants of lineage B.1.1.298. Cluster 4 was the most common virus variant found amongst these outbreaks (Figure 1) and was detected on 121 farms, while Cluster 2 and Cluster 3 viruses were found on 76 and 66 farms, respectively (note, some farms had viruses from more than one cluster present, see Supplementary Figure S1 and Discussion). In contrast, the Cluster 5 variant was only observed in mink from five farms in Northern Denmark (Table 1 and Figure 2A) and only during the first part of September 2020, whereas the other Clusters persisted until the culling of all mink in DK that ended in late November (Table 1). Further details of the various Clusters are described in Supplementary Information file S1.

The mink variant viruses with Y453F (within lineage B.1.1.298 including Clusters 2, 3, 4 and 5) clearly made up the majority of the variants found on Danish mink farms during the mink epidemic (Figure 1). However, new introductions of SARS-CoV-2 into mink also occurred, which lead to the C1-C7 variant groups (note a single virus sequence was assigned to C8, see Figure 1 and Table 1, but is not considered further). These new introductions occurred in multiple locations within Northern, Central and Southern Denmark (Figure 2B). These viruses are clearly distinct from the majority of those that infected the mink. For example, the viruses in C1-C7 lack the Y453F substitution in the S protein and they do not belong to the B.1.1.298 lineage. In total, mink on eighteen farms were infected with SARS-CoV-2 lineage variants other than B.1.1.298. These individual independent introductions are described in more detail in Supplementary Information file S2.

### Evolution of SARS-CoV-2 in mink and humans

In order to investigate the evolution of SARS-CoV-2 in mink and in humans within DK, the sequences of the viruses from both hosts during the same time period were compared. The full-genome sequences of SARS-CoV-2 from samples collected from Danish mink were collected from GISAID [21] and low-quality sequences (i.e. with more than 10 unresolved nucleotides) were removed. Sequences from humans in DK, circulating at that time, were also retrieved. For each of the datasets, identical or nearly identical sequences were removed (see Materials and Methods). The final data set comprised 258 sequences from mink on 129 farms and 497 sequences from humans across DK. These were aligned to the Wuhan-Hu-1 reference genome (GenBank accession no. NC_045512) as described, and a phylogenetic tree, including the mink and human viruses, was constructed (Figure 3). It is apparent that there was considerable heterogeneity among both the mink and human sequences in DK during this period. Furthermore, it can be seen that sequences derived from mink and human hosts are interspersed on the tree, indicating multiple cross-species transmission events occurred (Figure 3).

**Figure 3.**
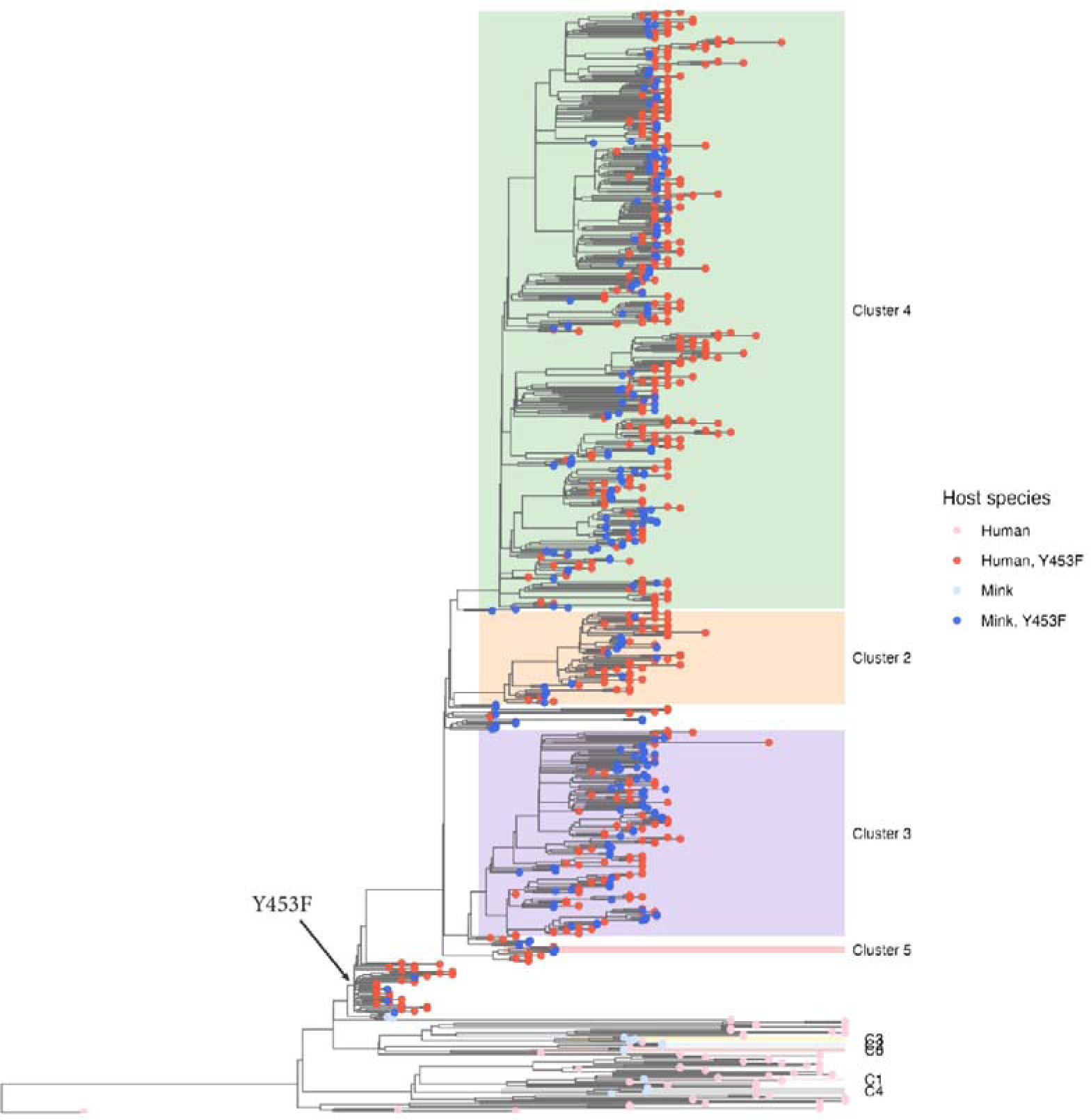
Phylogenetic tree based on whole-genome SARS-CoV-2 sequences from viruses obtained from humans and mink. Phylogenetic analysis was performed using BEAST2 with a strict clock model, GTR+gamma substitution model, and a BDSKY-serial tree prior. Shown here is the maximum clade-credibility (MCC) tree based on 17,500 post-burnin tree samples. Tips are colored based on host species (Human: red, Mink: blue), and on whether the encoded Spike protein contains the Y453F substitution (Yes: darker colors, No: lighter colors) resulting from the A22920T mutation. Cluster colors have been dimmed, compared to Figure 1, to emphasize the tip colors. The Y453F substitution can be seen to have evolved only once (arrow pointing to tree branch), after which point it was retained in all descendant viruses. Also note how mink and human sequences are interspersed indicating frequent cross-species jumps.

### Evolution and spread of mink-derived virus variants

At the time of the first introduction of SARS-CoV-2 into farm 1, in Northern Denmark (Figure 2A), the amino acid substitution Y453F, in the receptor-binding domain of the S protein (resulting from the mutation A22920T), had not been seen anywhere else (globally) except in mink from one of the infected mink farms in the NL. In this case, the substitution was in a different clade (19A) of SARS-CoV-2 [7, 8], so this finding did not indicate a connection between the outbreaks in DK and in the NL. Virus from the person connected to farm 1 in DK, who is presumed to be the source of the outbreak in mink, did not have this mutation in the spike protein gene. Indeed, the viruses from mink on farm 1 varied at this position, some had the A22920T mutation (resulting in the Y453F substitution) whereas others lacked this change [8] (Figure 1). Phylogenetic analysis based on whole-genome SARS-CoV-2 sequences from both mink and human hosts, also clearly showed that the Y453F substitution evolved only once (among mink on farm 1) and then spread, with all descendant mink- and human-derived sequences retaining this mutation (Figure 3 and 4).

**Figure 4.**
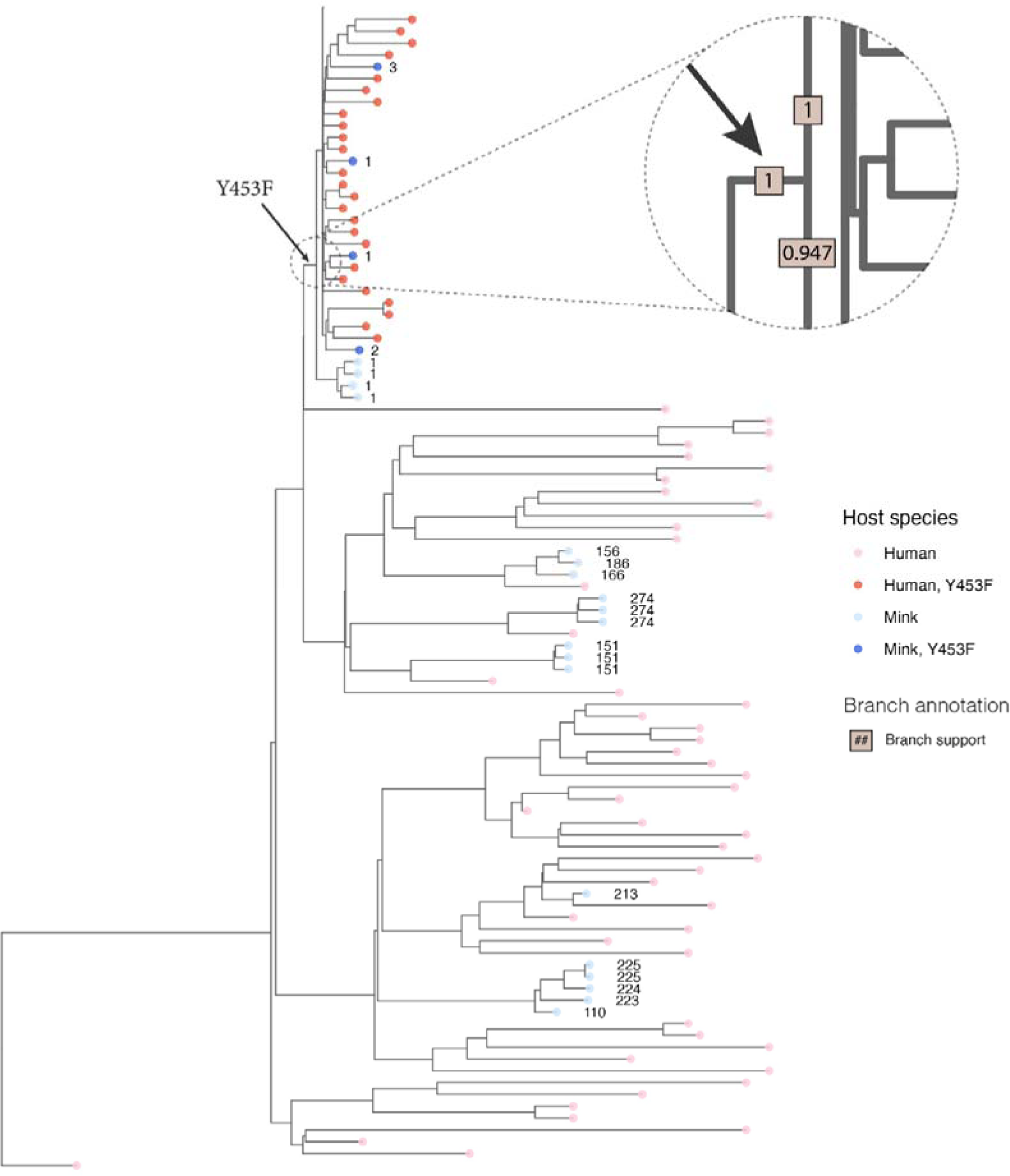
Zoom of phylogenetic tree from Figure 3 showing details around the branch where the Y453F substitution in the S protein occurred. Tips are colored based on host species (Human: red, Mink: blue) and on whether the encoded Spike protein contains the Y453F substitution (Yes: darker colors, no: lighter colors). Mink sequences are annotated with a number indicating the ID of the farm from which the sample was obtained. The dashed circle highlights the part of the tree where the Y453F substitution first appeared. The enlarged region is annotated with the posterior branch support values in brown boxes. Note how only farm 1 had some mink without the Y453F change (light blue) and some with it (dark blue). This is consistent with the substitution occurring in the mink on farm 1.

The deletion of residues H69/V70 in the S protein, on the other hand, appears to have evolved up to 5 times independently among the human and mink viruses analyzed here (Figures 5 and 6). One of these events occurred among the group of viruses in the mink that already had the Y453F substitution. The H69/V70del modification, as well as two other deletions in ORF1a, were observed for the first time on farm 4 [16]. Specifically, and based on the clock-tree reconstructed using BEAST2, the deletion resulting in the H69/V70del change evolved about 2-7 weeks after the appearance of the Y453F variant (Supplementary Figure S2). This is consistent with a previous analysis, which showed that deletion of H69/V70 from the S protein increases virus infectivity and compensates for an infectivity defect resulting from the RBD-substitutions N439K and Y453F [22]. All viruses, in the clade descending from this event, inherited this deletion, which was, therefore, present in the vast majority of the mink-derived viruses analyzed here.

**Figure 5.**
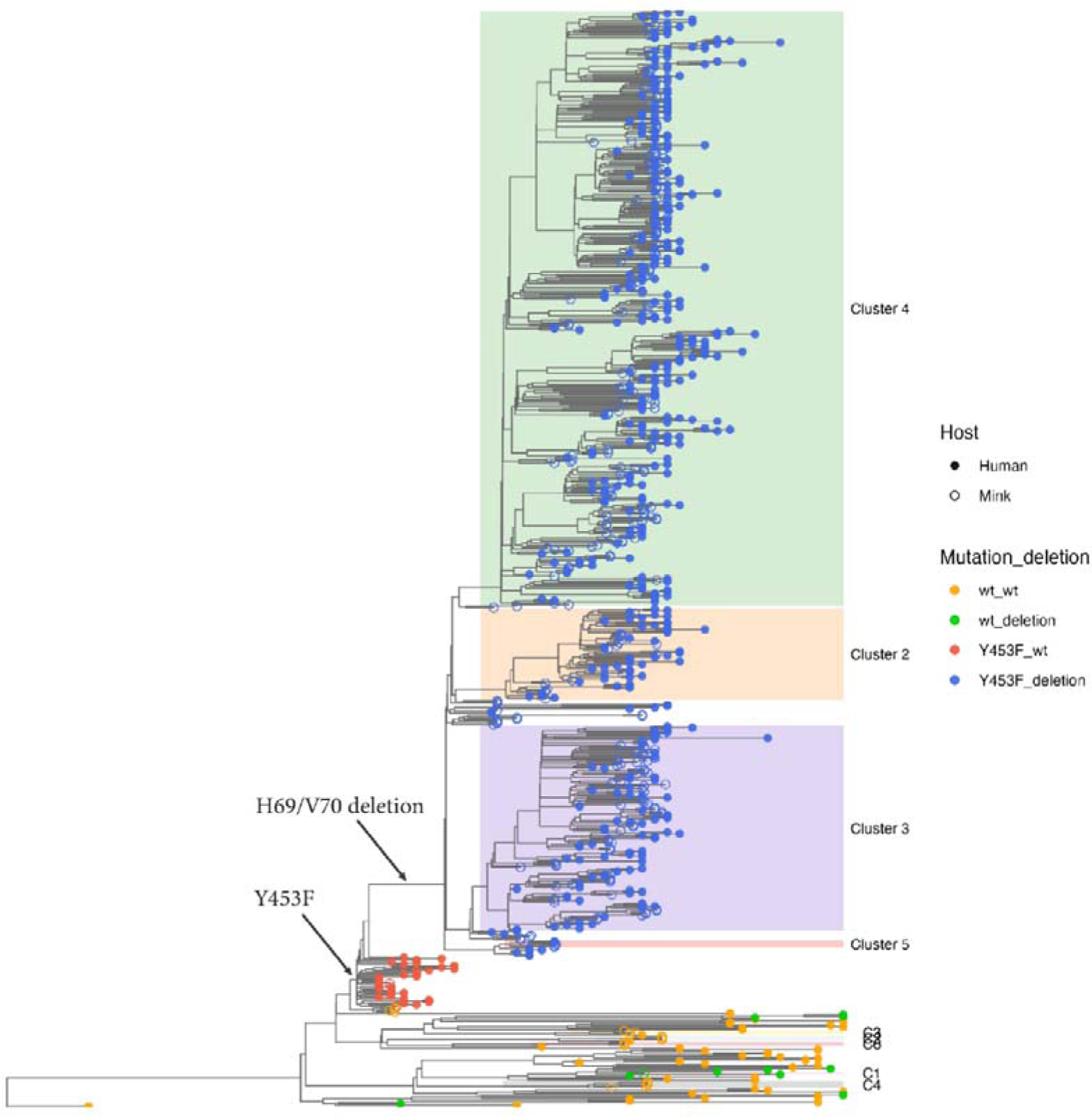
Phylogenetic tree from Figure 3 with tips colored according to presence or absence of the Y453F substitution and the H69/V70 deletion within the S protein. The format used to label tips is <Y453F status>_<DELETION status>, with “wt” indicating the absence of substitution or deletion, “Y453F” indicating the presence of that substitution, and “deletion” indicating the presence of the H69/V70 deletion: wt_wt: orange, wt_deletion: green, Y453F_wt: red, Y453F_deletion: blue. Host species is indicated using open circles for mink and closed circles for human. Note that the H69/V70 deletion appears shortly after the Y453F substitution (arrows pointing to branches), and both changes are subsequently present in all descendant sampled viruses, from both humans and mink. The deletion was also present in 4 separate clades among viruses without Y453F (4 groups of green tips in bottom part of tree – see Figure 6 for further detail).

**Figure 6.**
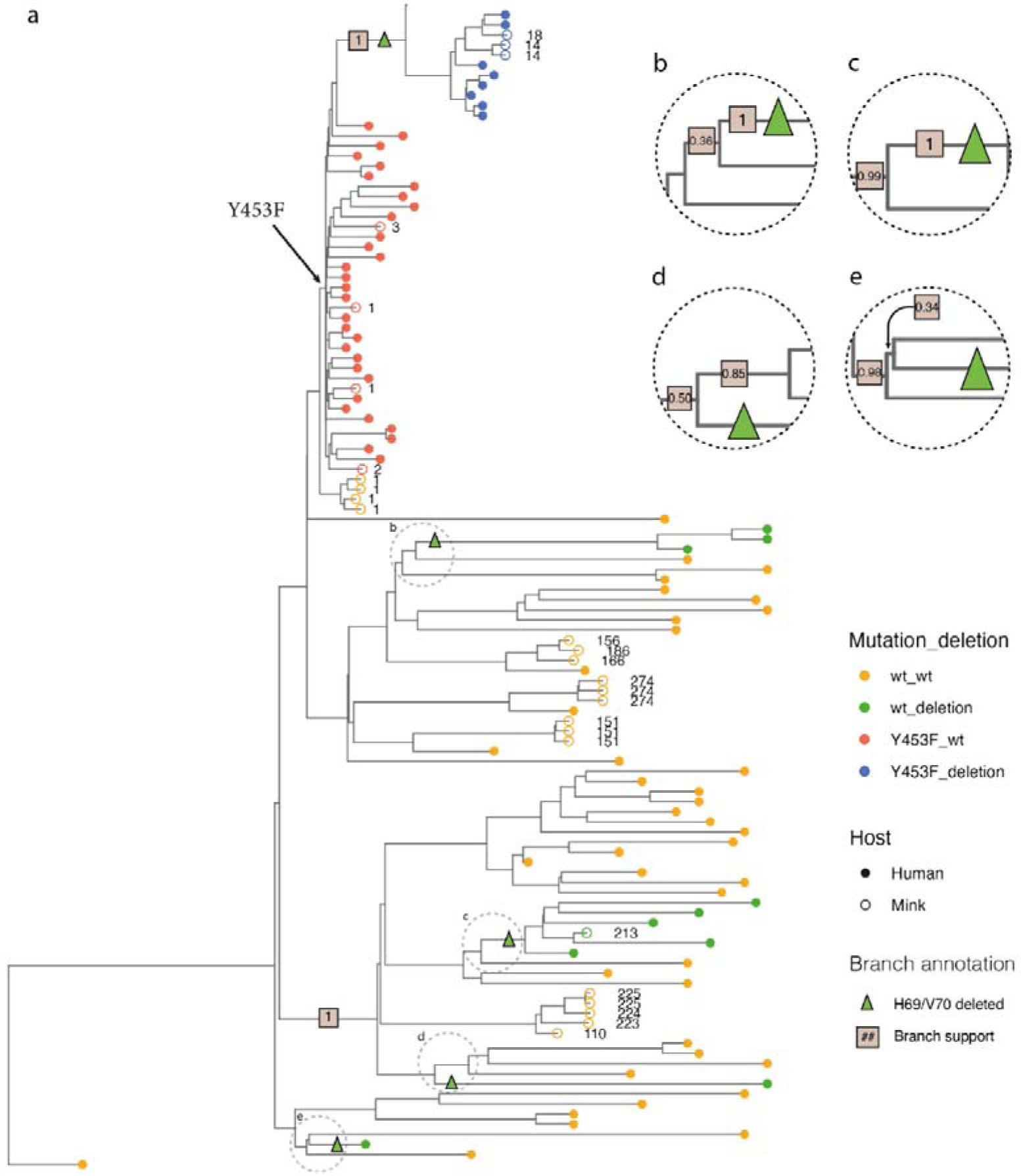
Zoom of phylogenetic tree from Figure 5 showing details around the branches where the H69/V70 deletion appeared. Color scheme is the same as in Figure 5. a) Mink derived sequences are further annotated with a number indicating the farm ID from which the sample was obtained. The deletion can be seen to have evolved on a branch shortly after the Y453F substitution and to then have been retained in all viruses descending from this branch (see upper part of tree in Figure 5). Among the viruses, that do not have Y453F, the deletion is present in 4 separate groups (see green tips in bottom part of tree). The basal branches (where the deletions presumably evolved) of these 4 groups are indicated with green triangles. A cutout has been made of each of these clusters (b-e) with annotated posterior branch support in boxes. Two of the 4 clusters are human singletons (green closed circles near bottom of plot) and may correspond to independent introductions into Denmark of viruses already harboring the deletion. The two other clusters contain multiple sequences (3 and 6 respectively), indicating that the deletion may have evolved in Denmark and subsequently spread. One of these clusters contains only human sequences, while the other contains the sequence from a single mink (from farm 213), that appears to have been infected by a human harboring virus with the deletion.

Among viruses, which do not have the Y453F substitution, the H69/V70 deletion appeared again in 4 separate locations on the phylogeny (Figures 5 and 6). Two of these are singleton human sequences, that are basal to the Danish sequences, and they may, therefore, represent separate introductions rather than cases where the deletion evolved among Danish viruses. In addition to these single leaves, there are two clades, within the non-Y453F part of the tree, where multiple related sequences all have the deletion (Figure 6). It appears that the deletion evolved independently among Danish viruses in these two cases, and then spread. One of these clades contains 3 human sequences, while the other contains 1 mink-sequence and 5 human sequences indicating that virus with the deletion was transferred between humans and mink. In some of these viruses, the H69/V70 deletion was coupled with the N439K substitution in the S protein, which is also within the RBD, and where the deletion has also been reported to function as a compensatory change [22].

### Inference of the number of cross-species transmissions in DK

We wanted to identify each of the cross-species transmission events, these could happen in either direction and both types of event are important for the spread of the infection. In previous studies, Wang et al. [23] defined criteria for identifying a cross-species transmission event for SARS-CoV-2 using a subset of the Danish sequences. These criteria were: (1) that the direct two branches after the root of the clade have a different host; and (2) that the posterior probability of both branch and ancestral host for the root of the clade is >0.8. In the dataset used by Wang et al. [23], three independent cross-species transmission events were observed, all of which were caused by human-to-mink transmission. In addition, six SARS-CoV-2 sequences from humans were found to be very similar to mink-derived viral genomes, indicating they were most likely transmitted from mink to humans. However, Wang et al. [23] concluded that they could not determine, using their analyses, how many independent cross-species transmission events occurred due to the low posterior probabilities of the branches.

In order to further investigate the incidence of cross-species virus transmission events, we have used the collected whole-genome sequences from DK (as described here) to infer the number of times that SARS-CoV-2 jumped from mink to humans (and *vice-versa*). Briefly, BEAST2 [24] was used to reconstruct clock model-based phylogenies. Then TransPhylo [25] was used to infer transmission trees based on the output from BEAST2, and finally the sumt and phylotreelib python packages [26, 27] were used to analyze the transmission trees and count the likely number of zoonotic and reverse zoonotic jumps between the two species. This number was calculated using four different methods (see Materials & Methods). In method A, the number of inferred direct transmissions from an observed mink sequence to an observed human sequence were counted. Using this approach, it was estimated that there had been about 9 direct transmissions (posterior mean: 8.6; 90% credible interval: 7-11) from one of the 258 mink sequences included in the dataset, to one of the 497 human sequences. In method B, *indirect* transmissions were also inferred from an observed mink sequence, via an unobserved intermediate host, to an observed human sequence. Using this approach, it was estimated that there had been about 17 jumps (posterior mean: 17.2, 90% credible interval: 14-20) from one of the mink to one of the humans in the data set. Using this same method, there were estimated to be about 18 jumps (posterior mean: 18.3; 90% credible interval: 15-21) from humans to mink. In method C, the number of cross-species jumps was estimated using a parsimony method applied to the TransPhylo output, including inferred unobserved mink and human hosts also. Using this approach, it was found that there had been about 59 jumps from mink to humans in DK during the investigated period (posterior mean: 59.0; 90% credible interval: 39-77). The result of method B, about 17 jumps from mink to humans, can be considered as a fairly high-confidence, but conservative, estimate, i.e., it is reasonably sure that the number of jumps is not less than this. However, since the virus from only a small proportion of the infected mink that were in DK during that time have been sequenced, it is almost certain that many interspecies jumps will be missed. The result from method C, i.e. about 59 jumps, may be argued to be probably closer to the real number as it represents a less conservative estimate. However, it comes with a greater uncertainty.

Using method C, a parsimony method applied to the TransPhylo output, it was estimated that there had also been about 136 jumps from humans to mink (posterior mean: 136.4., 90% credible interval: 117-164). This fits fairly well with the 129 different mink farms, with infected mink, represented in our data set, and is consistent with the hypothesis that most of the virus introductions into the mink farms have occurred by independent human-to-mink transmission events (not by mink from one farm directly infecting mink at another farm).

Finally, in method D we took an entirely different approach and instead used a structured coalescent, as implemented in the BEAST2 package MASCOT [28, 29]. Specifically, we fitted the model using two different sub-populations (mink and human) and hereby obtained the posterior distribution for the number of host state-changes in either direction. Using this approach we estimate about 30 mink-to-human host jumps (90% credible interval: 20-38) and 265 human-to-mink jumps (90% credible interval: 248-281). These estimates are in good agreement with the values found using the methods B and C.

The different methods mentioned above have different strengths and shortcomings, and account for different aspects of the underlying processes (see Discussion), but it is reassuring that they nevertheless give qualitatively similar results (see Table 2).

**Table 2.** showing number of cross-species jumps inferred using different approaches (posterior mean and [90% highest density credible interval])

| Method | Details | Mink to Human | Human to Mink |
| --- | --- | --- | --- |
| Method A | Transphylo: direct transmission between two observed sequences | 8.6 [7, 11] | 13.2 [10, 15] |
| Method B | Transphylo: direct + indirect transmission between two observed sequences; only unique infecting hosts are counted. | 17.2 [14, 20] | 18.3 [15, 21] |
| Method C | Jumps counted on transphylo transmission tree with all unobserved node states inferred by parsimony | 59.0 [39, 77] | 136.4 [117, 164] |
| Method D | BEAST2 + MASCOT: structured coalescent | 30.1 [20, 38] | 265.1 [248, 281] |

## Discussion

SARS-CoV-2 infection of farmed mink in DK contributed to the epidemic in humans in DK during 2020. The epidemic in mink was not being efficiently controlled by the measures taken (mink on 290 farms out of about 1100 in the country were found to have been infected) and it was decided to cull over 15 million mink. This resulted in the closure of the mink production industry until after the end of 2022. Most of the outbreaks in mink were caused by one of three different virus lineages, termed Clusters 2, 3 and 4, all of which belong to the pangolin lineage B.1.1.298 (Figure 1). These clusters shared some common features, namely the H69/V70del and Y453F changes, within the S protein. The deletion of H69/V70 has arisen independently in a variety of different lineages of SARS-CoV-2, both within mink and human variants. The deletion is associated with increased cleavage of the S protein and confers enhanced virus infectivity [22].

A virus isolate from Cluster 5, with additional amino acid changes, was the focus of considerable attention since preliminary studies indicated this isolate showed resistance to neutralization by antibodies from a small panel of convalescent human patients [18]. However, in follow up studies [19], it was found that the antibodies from just 3 out of 44 patient samples tested had a >3-fold reduction in virus neutralization titer against the Cluster 5 virus isolate compared to a virus from early in the pandemic. Only one sample from the 44 patients had a neutralization titer that was reduced by 4-fold or more [19]; the latter being the threshold set for defining neutralization resistance [30].

The Y453F substitution was found to have evolved only once in the mink in DK, on farm 1 [8]. This change was present in the majority of the sampled mink sequences (Figure 1 and Suppl. Figure S1) and was also found in sequences from more than 1100 human cases in DK. It has been estimated that about 4000 humans have been infected with this variant [11]. Thus the Y453F change clearly does not have a severely detrimental effect on the ability of the virus to infect humans [31]. However, viruses with this change were rapidly lost following the culling of all the mink (Table 1 and [11]). Cluster 5 viruses were not detected in mink or humans after mid-Sept. 2020 but viruses of the B.1.1.298 lineage (with the Y453F change) were detected in humans until January 2021 [17]. This suggests that viruses with the Y453F change did not have a selective advantage in humans at this time point. In principle, it could have been possible that the Y453F variant was outcompeted by the arrival of another variant in the human population. Indeed, the Alpha variant of SARS-CoV-2 was first detected in Denmark at the end of 2020, and later became dominant, but it only represented a small fraction of the infections in the population until well after January 2021 by which time the Y453F variant had already disappeared (see https://www.covid19genomics.dk/statistics). Thus, it does not seem possible to attribute the loss of the Y453F variant to competition from the Alpha variant in the human population. It is noteworthy that the generation of the Y453F variant (with the H69/V70del) in a human patient with lymphoma has been reported [32], in a virus lineage separate from the mink viruses. As indicated above, the Y453F change only occurred once in mink in DK, however, it is notable that this change also has occurred independently in other mink virus sequences in the NL [7], Poland [33], the USA [34] and (based on sequences from GISAID [21]) in Lithuania and Latvia. All of these changes occurred in lineages other than B.1.1.298, indicating convergent evolution due to selective advantages in mink. It should be noted that all but one sequence within the B.1.1.298 lineage originated from DK [21]. The single sequence from outside DK was found in a human sample collected in the Faroe Islands (these have close links to DK) in September 2020.

In the lineage C4, which was first recognized in mid-October 2020 (i.e. shortly before the cull commenced) and lacks the Y453F change, another change, N501T, was detected on multiple farms (Supplementary Information file S2). Like the Y453F change, this substitution occurs at the interface between the ACE2 receptor and the S protein. Thus, it may achieve a similar effect [31]. It is notable that this change has also occurred in mink sequences from multiple countries and in different virus lineages as for the Y453F substitution (see above).

Assessing the extent of interspecies virus transmission is not simple, see Wang et al. [23]. Due to the many highly similar sequences, there will be several branches in the phylogenetic tree with poor support, and this causes what may be termed an entropic problem leading to an upward bias in the count of interspecies jumps [35]. If a set of, say, 5 mink sequences and 5 human sequences each have one unique mutation, then their pairwise distances will all be 2, and all the possible resolutions of this 10-leaf subtree will be equally likely. However, since there are many more possible subtrees where the 5 mink and 5 human leaves are intermingled, than there are possible subtrees where they are cleanly separated, then the average number of inferred jumps will be biased towards more than 1 inter-species jump, even though the data would also be consistent with only one zoonotic event. This means that ordinarily used methods for dealing with phylogenetic uncertainty, such as performing the computation on all or many trees from BEAST’s posterior sample, will not work (instead of getting a reliable posterior count, accounting for the uncertainty, the inclusion of less supported trees will create a bias for over-counting).

Here, we have used four different methods to assess interspecies virus transmission. Using the method B, the analysis of the sequences indicated that about 17 (90% credible interval: 14-20) different mink to human transmission events have happened in DK. This was estimated using a very conservative approach. Using an alternative method, based on analyzing the output from TransPhylo using parsimony (termed here method C), about 59 jumps from mink to humans were estimated to have occurred. Furthermore, this methodology generated an estimate of 136 jumps from humans to mink. This number fits well with the 129 farms represented in the data set that had infected mink. Using an entirely different method, based on a structured coalescent model, we estimate about 30 mink-to-human and 265 human-to-mink cross-species jumps. Given the very different set of assumptions for these models we find the above estimates to be qualitatively in good agreement (see Table 2).

It should be noted that the two main approaches (using TransPhylo or using MASCOT) have different strengths and weaknesses and account for different aspects of the underlying processes to varying degrees. Thus, the approach where we used BEAST+TransPhylo explicitly takes into account the difference between a phylogeny on the one hand (where nodes correspond to viral particles, and branches correspond to replication events) and a transmission tree on the other hand (where nodes are host organisms, and branches are transmission events, and where there can be viral diversity within a host). This distinction is not made when using MASCOT. Additionally, TransPhylo explicitly uses the information about sampling dates and the typical duration of being infectious, when computing the likelihood of a transmission tree (two sequences sampled 7 days apart are more likely to be part of a transmission event, than two sequences sampled 40 days apart, for instance). Again, MASCOT cannot use this information when estimating the number of state changes (= cross-species transmission events).

On the other hand, a structured coalescent model like MASCOT could be a good description of many of the important aspects of what is really happening here, and especially so if we have one sub-population for each farm. On any given farm the infection is likely to spread rapidly and R0 is likely very high compared to the R0 among humans. It would make sense to explicitly account for this difference in the model parameters. However, it is computationally essentially impossible to have one sub-population per farm: even with the marginalization and other approximations used in the MASCOT package, this approach still requires solving a number of differential equations that is proportional to the number of states times the number of lineages. This is computationally unfeasible with > 100 states (individual farms) and hundreds of lineages. One alternative is then to have only two states: mink and human. However, in this case the structured coalescent model assumptions will fit very poorly to reality. On the whole, mink from one farm never interact with mink from a different farm. At the same time, we typically only have 1, 2, or 3 sequences from each farm, while having 258 mink sequences in our entire data set. That means that there is a very small probability that any given mink lineage will coalesce with any other mink lineage (the two lineages are most likely to be on two different farms), and a much higher probability that a mink lineage will coalesce with a human lineage. It is, then, not entirely clear what the coalescence rate within the mink subpopulation will actually mean when estimated from data such as these (and it certainly should not be expected to be informative about the rapid spread inside mink farms).

Another potential issue with our analyses is how well we account for the phylogenetic uncertainty, given that we use a quite limited number of trees from the posterior tree samples in the BEAST2 output. Thus it might be the case that including more BEAST trees would lead to significantly wider credible intervals for the estimated number of cross-species jumps (less certainty about our estimates). To test whether this was the case we performed an analysis on a subset of about 200 sequences where we compared the width of credible intervals when using 100 BEAST trees as input to TransPhylo, or only 25 BEAST trees (data not shown). The main conclusion from this analysis is that we get essentially the same estimated uncertainty (width of CI) when using 25 as when using 100 BEAST trees, and we therefore think this issue has a limited impact on our results.

The transmission of the mink variant viruses from one mink farm to another occurred very frequently. However, the mechanisms involved in this spread are not established [9]. In previous analyses, there has been shown to be no significant support for transmission from one farm to another by wildlife (including birds), feed suppliers or visiting veterinarians [9]. Only proximity to an infected farm and the size of the farm (and hence the number of staff) were associated with the risk of animals on a farm becoming infected. The results presented here are consistent with this information. Thus, it seems that most of the introductions of the virus onto these mink farms have occurred by independent mink-to-human and then human-to-mink transmission events (not by mink from one farm directly infecting mink at another farm). Airborne transmission of the virus from mink farms to humans not connected to the farm seems unlikely, since the concentration of virus in the air outside of the mink farms appears to be low [9]. However, this topic deserves further study. One relevant observation is that, in the majority of cases, mink sequences from a single farm cluster together, suggesting that there has been a single introduction on each of these farms. Specifically, we have more than 1 mink sequence from 82 of the 129 farms analyzed here. On 8 of these farms (with farm IDs: 13, 27, 117, 121, 144, 148, 273, 279), the sequences belong to two different clusters, indicating >= 2 separate introductions. Thus, we can say that in 90% of cases (76/82), where we have data to address the question, there is a single observed introduction. Furthermore, from among all the 266 mink farms represented, multiple sequences were found on 214 farms. From these, there were 21 farms with sequences from 2 (for 20 farms) or 3 (for 1 farm) clusters, indicating 2 or 3 separate introductions onto these farms.

The major proportion of the viruses that infected mink in DK had the Y453F substitution together with the H69/V70del in the S protein, including all of the viruses in Clusters 2, 3, 4 and 5 (Figure 1). This suggests that, although new introductions of the virus, not derived from mink but from humans, occurred (as with C1-C7), these were much less important for the total outbreak in mink than the mink farm to mink farm transmission. It is clearly not possible to know whether some of these virus variants would have become predominant among the mink if they had not been culled.

### Concluding remarks

It is apparent that SARS-CoV-2 readily infected farmed mink and spread frequently between farms. Transmission from infected humans to mink and from infected mink to humans occurred on multiple occasions and the mink-derived viruses then spread among people. Movement of the mink-derived viruses between the mink farms is consistent with transmission via infected people. There were legitimate concerns that replication of SARS-CoV-2 in a large population of mink could generate novel variants that would have adverse effects on human health due to antigenic change, greater transmissibility or higher fitness. However, mink-derived viruses with such unwelcome characteristics did not spread among humans before the mink population was culled. Variants of SARS-CoV-2 that did arise in mink (e.g. with the changes Y453F and H69/V70del in the S protein) were transmitted to, and within, the human population but died out either before, or soon after, the culling of the mink population in DK.

## Materials and Methods

### Mink sample collection

Throat swab samples were collected from mink and analyzed as described previously [8].

### Map creation

The maps were made using the open source geographic information system Q-GIS with the shapefile of the administrative units of Denmark uploaded from DIVA-GIS (open source), onto which the data were plotted, see https://www.qgis.org/en/site/getinvolved/governance/trademark/index.html

### Sequencing strategy

Whole genome amplification of SARS-CoV-2 in mink and human samples was performed using a modified ARTIC tiled PCR protocol (see [36]) with amplicons ranging from 1000-1500 bp. A custom 2-step PCR with barcoding was applied to the amplicon libraries, then the libraries were normalized, pooled, and sequenced using Oxford Nanopore’s SQK-LSK109 ligation kit on a MinION device with R.9.4.1 flowcells. The full protocol is available [37].

### Construction of maximum likelihood phylogenetic tree

The maximum likelihood phylogeny of all 698 SARS-CoV-2 sequences from mink isolates was reconstructed using IQ-TREE version 2.0.3 [38] with a GTR model, based on the alignment obtained by comparing each sequence to the Wuhan-Hu-1 reference genome (GenBank accession no. NC_045512) using MAFFT version 7.475 [39] with option ‘--addfragments’. A list of mink derived SARS-CoV-2 sequences from Denmark is available with the GISAID identifier EPI_SET_240529zr, (available at https://doi.org/10.55876/gis8.240529zr).

The phylogenetic tree was thereafter annotated using package ggtree in R version 4.2.1 [40]. Clusters 2-5 were derived from the initial cases (on farms 1-5) while the separate introductions that resulted in the C1-C8 variant groups were defined from a phylogeny based on human and mink sequences by picking the smallest possible monophyletic group containing one or more mink sequences.

### Construction of Bayesian phylogenetic trees

Whole-genome sequences of SARS-CoV-2 derived from infected farmed mink and humans in DK were collected from GISAID [21] on August 31^st^ 2023. Sequences derived from mink were collected by searching for complete sequences passing GISAID’s high coverage filter (allowing only entries with <1% Ns and <0.05% unique AA mutations) with a precise collection date. These gave rise to dataset 1, a subset of the dataset presented above recreated from public sources. The sequences downloaded can be found using the EPI_SET identifier EPI_SET_240506tk, available at https://doi.org/10.55876/gis8.240506tk. For this dataset, consisting of mink virus sequences, duplicate sequences derived from samples from the same farm on the same date were removed. Similarly, sequences derived from humans were collected by searching for complete sequences with a collection date between June 1^st^ 2020 and February 28^th^ 2021 passing GISAID’s high coverage filter. The sequences downloaded can be found using the EPI_SET identifier EPI_SET_240506yv, available at https://doi.org/10.55876/gis8.240506yv. Two different datasets were constructed consisting of human virus sequences: dataset 2 with the amino acid substitution S:Y453F and dataset 3 without the amino acid substitution S:Y453F. For datasets 2 and 3, duplicate sequences were removed if they were sampled on the same day. This was done to preserve the temporal signal in the data.

Sequences with more than 10 undetermined nucleotides were removed from the datasets, and the datasets were pre-processed by masking as described [41], removing sequences with more than 100 end gaps. Dataset 3 was further reduced to minimize the computational load using CD-HIT-EST from CD-HIT [42] to achieve a representative dataset using a similarity threshold of 0.999. The three datasets were combined into one consisting of 258 sequences from mink (derived from 129 mink farms), 49 sequences from humans without the S:Y453F substitution and 448 sequences from humans with the S:Y453F substitution. These sequences were aligned as described above. We here note that subsampling using CD-HIT was performed only on human sequences not encoding the Y453F substitution. However, the vast majority of cross-species transmissions appear to occur in the other part of the tree (the one with the Y453F substitution), where subsampling of human sequences was not performed. The impact of subsampling of human sequences on the estimated number of cross-species transmission events, is therefore expected to be low.

### Estimation of the number of zoonotic jumps from mink to human

To determine transmission pathways, information from the phylogenies together with the relative sampling dates was combined. Phylogenetic trees were reconstructed using BEAST 2 [24]. The substitution model was GTR with empirical base frequencies and gamma-distributed rates with 4 discrete categories, combined with a strict molecular clock model calibrated by using the sequence sampling-dates, obtained from GISAID, to date the tips of the tree. The tree prior was the birth-death skyline serial model, with 10 dimensions for the reproductive number parameter, and one dimension for the sampling proportion [43]. The model estimates a separate effective reproduction number for each of 10 equally large time-intervals covering the time-span from the root of the tree to the farthest tip. The prior for the becoming-uninfectious rate parameter was lognormal(M=52.0, S=1.25, mean in real space) per year, corresponding to a prior 95% credible interval of [1.3, 180] days for the duration of an infectious period. The prior for the clockrate was lognormal(M=0.001, S=1.25, mean in real space) substitutions per site per year, corresponding to a 95% prior interval of [4.0E-5, 5.3E-3] substitutions per site per year. Both of these priors are weakly informative and help to regularize model fitting without imposing very strict constraints on the estimated values for these parameters. Other priors were left at their default values. Two parallel MCMC chains were run for 50 million iterations each with logging of trees and other parameters every 4000 iterations (for a total of 2 x 12,500 parameter samples). A burn-in of 30% (15 million generations) was used. The software Tracer v1.7.2 [44] was used to analyze parameter samples. Marginal posterior distributions from the two runs were essentially identical, indicating good convergence. Effective sample sizes for all parameters were well above 200, except for the following: posterior (ESS=166), likelihood (ESS=94), tree-length (ESS=136), BDSKY_serial (ESS=138). The software phylotreelib [26] and sumt [27] were used to analyze tree-samples, and to extract post-burnin trees and compute maximum clade credibility trees. Tree samples from the two independent runs were very similar, with average standard deviation of split frequencies (ASDSF) of 0.0125. The number of effective tree samples was estimated by first computing the log clade credibility for each tree-sample (based on clade frequencies from all post-burnin trees), and then using Tracer to compute ESS from this proxy measure [45]. Computed this way, the tree-sample ESS was 287, indicating an acceptable number of independent tree samples in the posterior.

To infer transmission trees, the software TransPhylo v1.4.10 [25] was used. This takes as input a pre-computed, dated phylogeny, where leaves correspond to pathogens sampled from the known infected hosts. The main output is a transmission tree that indicates “who” infected “whom”, including the potential existence of unsampled individuals who may have acted as missing transmission intermediates. For input we used the maximum clade credibility (MCC) tree with common-ancestor depths. A further 28 randomly selected other trees from BEAST2’s posterior samples were analyzed, chosen to cover a range of different log-clade credibility values. We also used common-ancestor depths to set the branch lengths of these trees. Before analyzing any of these trees, the original Wuhan sequence was removed from the tree with the aim of having a more homogeneous substitution process on the remaining branches for the TransPhylo analysis. The generation time distribution in TransPhylo was set to be gamma-distributed with shape-parameter=60 and scale-parameter=0.0004105. These parameters were chosen to match the posterior 95% credible interval, found in the BEAST-analysis, as closely as possible (6.86 to 11.4 days). The parameters were found using the optimize.minimize function from the SciPy python package [46]. TransPhylo was run for 10 million iterations, sampling every 2000 generations, and using a burnin of 50%. This gave a total of 2500 post-burnin samples of transmission trees and other parameters, for the MCC tree and for each of the 28 other trees from the BEAST posterior sample. For the TransPhylo run, we set updateOff.p=TRUE to allow estimation of the offspring distribution. Convergence was checked by inspecting trace plots and computing ESS.

The output from TransPhylo was further analyzed to estimate the number of times SARS-CoV-2 jumped between mink and humans. This was done by inspecting each of the 72,500 posterior transmission-tree samples (i.e. 29 times 2500), and for each of them counting the number of jumps in three different ways. In method A: the inferred direct transmissions from an observed mink sequence to an observed human sequence were counted (i.e., cases where TransPhylo inferred that both the source and the target of a cross-species transmission event were included in the data set). In method B: the number of inferred *indirect* transmissions from an observed mink sequence to an observed human sequence were counted. Occasionally TransPhylo will infer transmission chains that include one or more unobserved links (e.g., mink -> unknown -> unknown -> human), and these, of course, also imply transmission of the virus from a mink to a human somewhere in that chain. In method C: a parsimony method was used to infer the minimum number of mink-to-human transmissions based on the posterior sample of the transmission trees inferred by TransPhylo. Specifically, the algorithm of Hartigan [47] was implemented in a version that allowed some internal nodes on the tree to be observed (i.e., their state sets are simply taken to be the observed host for that internal node).

An additional method employed for estimating the number of zoonotic jumps, D, was based on a structured coalescent model as implemented in the BEAST package MASCOT [28, 29]. We ran two parallel MASCOT analyses for 30M generations, with an HKY substitution model and with two sub-populations (mink and human). A burn-in of 33% (10M generations) was used. Convergence was checked as described for BEAST above. Effective sample sizes for the following parameters were below 200: posterior (ESS=102), likelihood (ESS=149), prior (ESS=164), treeLikelihood (ESS=149), Tree.treeLength (ESS=116), clockRate (ESS=149), Mascot (ESS=173).

### Pangolin lineage determination

The Pangolin lineage for the individual variants has been determined by analysis of the mink sequences in the database PANGO lineages [48]. In addition, a search in the GISAID EpiCoV database [21] has been used for further analysis in order to examine the occurrence of selected variants in published mink sequences and human sequences. When describing the observed changes in the S protein, the change D614G (compared to the reference Wuhan strain) was omitted, as this change occurred very early in the pandemic and is present in all sequences during the period of interest.

## Author contributions

Conceptualization: T.B.R., A.Bø., G.J.B.; Data curation: T.B.R., A.S.O.; Formal analysis: T.B.R., A.G.Q, A.G.P., M.J.F.G., E.R.T., J.F., M.R.; Funding acquisition: A.Bø., T.B.R., G.J.B., A.G.P.; Investigation: T.B.R., R.N.S., M.S., L.L. A.Bo.; Methodology: R.N.S., M.S., A.G.Q., M.J.F.G., E.R.T., A.G.P.; Resources: L.L., S.M. (study materials); Software: R.N.S., A.G.P., A.G,Q, M.J.F.G., E.R.T.; Supervision: T.B.R., A.G.P., A.Bø.; Validation: T.B.R.; Visualization: T.B.R., A.G.Q., F.F.C.A., M.J.F.G., E.R.T., R.N.S., A.G.P., A.Bo.; Writing-original draft preparation: G.J.B; Writing-review & editing: All authors.

## Acknowledgements

We gratefully acknowledge all data contributors, i.e., the Authors and their Originating laboratories responsible for obtaining the specimens, and their Submitting laboratories for generating the genetic sequence and metadata and sharing via the GISAID Initiative, on which this research is based. We thank Kåre Mølbak for helpful comments on an early draft of the manuscript.

This research was funded by the Danish Veterinary and Food Administration (FVST) as part of the agreement for commissioned work between the Danish Ministry of Food and Agriculture and Fisheries and the University of Copenhagen and the Statens Serum Institut. Further funding has been received from the Danish National Research Foundation (grant number DNRF170 to A.G.P).

## Supporting information

**Rasmussen et al. Mink Phylogeny Supplementary Information S1.**

Description of the major mink virus Clusters, termed 2, 3, 4 and 5

**Supplementary information on mink virus variants S2.**

Mink virus variants C1-C8 that lacked the Y435F substitution (additional introductions).

**RasmussenetalMinkphyloSupplementaryFigure S1.**

Maximum-likelihood phylogenetic tree generated using 698 mink derived virus sequences. The sampling dates and the farm identifiers are also indicated. The Clusters 2-5 (all with the Y453F substitution) are marked as are the C1-C7 variants which lack this change.

**Rasmussen et al., Mink phylogeny Supplementary Figure S2.**

Colored phylogeny, showing inferred chain of SARS-CoV-2 transmission from a mink (sequence*-*name shaded in blue) to humans (names shaded in red). The phylogeny shown here is a detail of a larger inferred transmission tree inferred by the software TransPhylo [25]. Different branch colors correspond to different hosts, observed or un-observed, and a region of connected branches with a single color represents evolution happening inside a single host. Stars on branches correspond to transmission events. If a colored branch is connected to a leaf, then the host has been observed, otherwise it is an un-observed intermediate host inferred by TransPhylo. The mink host (branches colored blue, name shaded in blue) was sampled on October 10, and is inferred to have transmitted the virus to two humans, that were both sampled on October 26 (branches colored orange and brown). One of these human hosts was inferred to further have transmitted the virus to another human, sampled on November 2 (branch colored grey, sequence name: 2020-11-02_H_670704).

