## Supplementary information S1 for "Emergence and spread of SARS-CoV-2 variants from farmed mink to humans and back during the epidemic in Denmark, June-November 2020"

**Rasmussen et al. Mink Phylogeny Supplementary Information** **S1**

*Description of the major mink virus Clusters, termed 2, 3, 4 and 5*

a) *Cluster 2*

Sequences belonging to Cluster 2 were found for the first time in samples taken on 9^th^ Sept. 2020. In total, 174 sequences have been found within Cluster 2 distributed across a total of 76 mink farms. The majority of these sequences (134 out of 174) did not encode additional amino acid changes, compared to the index case on farm 1, in the S protein (apart from H69/V70del and Y453F) but several changes outside of the S protein coding region were found compared to the reference strain (from Wuhan). However, in the remaining 40 sequences, from a total of 17 mink farms, further changes were found in the S protein (Table S2). The N751Y change was found for the first time on farm 61, in samples taken on 6^th^ Oct. 2020, and was later found on a further 15 farms. In three of these, it was seen together with A626S, while in eight mink farms it was found together with two further changes in the S protein (L5F and C1250F).

| No. of farms in Cluster 2 | No. of sequences | New S-protein substitutions |
| --- | --- | --- |
| 59 | 134 | - |
| 1 | 2 | D1139Y |
| 5 | 11 | N751Y |
| 3 | 6 | N751Y, A626S |
| 8 | 21 | N751Y, L5F, C1250F |

**Table S2.** Additional amino acid substitutions within the S protein of Cluster 2 viruses compared to the index case.

b) *Cluster 3*

Sequences belonging to Cluster 3 were found for the first time in samples taken from farm 15 on 14^th^ Sept. 2020. The farm 15 sequences are very similar to those found in samples taken from farm 5 on 26^th^ Aug. 2020 and it seems that farm 5 sequences are parental to Cluster 3 (see Supplementary Figure S1). In total, 142 sequences comprise Cluster 3 that originated from 65 mink farms. Of the 142 sequences, 50 had no additional changes in the S protein beyond H69/V70del and Y453F but several changes outside of the S protein coding region were found compared to the index strain. However, in 92 sequences from a total of 44 mink farms, further changes were found in the S protein sequence (see Table S3). The majority of these (86 sequences from 39 mink farms) only had the change S1147L, which was found for the first time on farm 31 in the sample taken on 25^th^ Sept. 2020. However, another five farms had viruses with an additional, distinct, single amino acid change in the S protein (Table S3).

| No. of farms in Cluster 3 | No. of sequences | New S protein substitutions |
| --- | --- | --- |
| 24 | 50 | - |
| 37 | 86 | S1147L |
| 1 | 1 | S1147L, Q675H |
| 1 | 2 | S1147L, Q836E |
| 1 | 1 | S1147L, Q613H |
| 1 | 1 | S1147L, T573I |
| 1 | 1 | S1147L, A222V |

**Table S3.** Additional amino acid substitutions within the S protein of Cluster 3 viruses compared to the index case.

*c) Cluster 4*

Genomic sequences located within Cluster 4 in the phylogeny are very closely related to those from the early cases from August (farm 4, see Supplementary Figure S1). The Cluster 4 viruses were detected for the first time in samples taken from farm 13, on 10^th^ Sept. 2020. In total, 272 genomic sequences belong to Cluster 4 and were distributed across 121 mink farms. Among these viruses, 247 sequences from 110 mink farms did not have changes (compared to the index case) in the S protein apart from the H69/V70del and Y453F. However, in 25 sequences from a total of 12 mink farms, further single amino acid changes have been found in the S protein (Table S4), most of these were only identified on a single farm. Several nucleotide changes, outside of the S protein coding region, were identified compared to the index strain.

| No. of farms in Cluster 4 | No. of sequences | New S protein substitutions |
| --- | --- | --- |
| 109 | 247 | - |
| 1 | 1 | G446V |
| 1 | 3 | K77N |
| 2 | 5 | V213E |
| 1 | 2 | D574Y |
| 1 | 1 | A684S |
| 1 | 1 | G1085V |
| 1 | 6 | S939F |
| 1 | 1 | T912I |
| 1 | 3 | V1264F |
| 1 | 1 | P9L |
| 1 | 1 | W64L |

**Table S4.** Additional amino acid substitutions within the S protein of Cluster 4 viruses compared to the index case.

*d) Cluster 5*

Cluster 5 consisted of virus sequences from mink on farms 6, 9, 10, 14 and 18, which were found positive for SARS-CoV-2 in the period from 31^st^ Aug. 2020 to 15^th^ Sept. 2020 (see Table 1). In addition to the Y453F change and the deletion H69/V70, the sequences within Cluster 5 also encode the amino acid substitutions I692V and M1229I in the S protein, as well as a variety of other changes elsewhere within the genome, as described previously [19]. As indicated above, preliminary observations indicated that a virus isolate belonging to Cluster 5 was antigenically distinct from the parental strain [18]. Viruses of this type were not found on any further mink farms but were found in 12 human cases in northern Jutland from August to September 2020 [11].
