## Supplementary information S2 for "Emergence and spread of SARS-CoV-2 variants from farmed mink to humans and back during the epidemic in Denmark, June-November 2020"

**Supplementary information on mink virus variants S2.**

*Mink virus variants C1-C8 that lacked the Y435F substitution (additional introductions)*

The mink variant viruses with Y453F (within lineage B.1.1.298 including Clusters 2, 3, 4 and 5) made up the majority of the variants found on Danish mink farms during the mink epidemic (Figure 1). However, new introductions of SARS-CoV-2 into mink also occurred, which lead to the C1-C8 variant groups (Figure 1). These new introductions occurred in multiple locations within Jutland (Figure 2B). These viruses are clearly distinct from the majority of those that infected the mink. For example, the viruses in C1-C8 lack the Y453F substitution in the S protein and they do not belong to the B.1.1.298 lineage. The first of these variants were from samples collected on 16^th^ Oct. 2020 on farms 110 and 111 and in samples taken on 26^th^ Oct. 2020 from farm 166 (Table 1). Subsequently, three more independent introductions, presumably from humans, have been found. Thus, in total, there were seven groups of variants, each lacking the Y453F change in the S protein, which are named: C1, C2, C3, C4, C6, C7 and C8 (note, C5 was omitted to avoid possible confusion with the widely discussed Cluster 5). During the last phase of the mink epidemic, some of these additional lineages of SARS-CoV-2 were detected from single farms, e.g. B.1.258.9 (C1) and B.1.1.159 (C7), but others, e.g. B1.1.219 (C2) and B.1.536 (C4), were found on three to seven farms (see Table 1). In total, mink on eighteen farms were infected with SARS-CoV-2 lineage variants other than B.1.1.298. These individual independent introductions are described in more detail below.

*a) C1 variant*

The C1 variant was first found in samples from mink taken on 3^rd^ Nov. 2020 from farm 213 in southern Denmark (Figure 2B). Mink sequences from this farm belong to the pangolin lineage B.1.258.9, which has lineage-defining changes that encode the following changes within the S protein: H69/V70del, N439K and G1223S (compared to the index strain). Although the H69/V70del change corresponds to the same deletion found in the mink variants with Y453F from Clusters 2 to 5, the C1 variant is not linked to these. The variants with the N439K substitution were found during November 2020 in about 10% of the sequenced human samples in DK [17]. There were 198 sequences from humans belonging to the B.1.258.9 lineage during the period 31^st^ Aug. 2020 to 25^th^ Jan. 2021. The C1 variant was not found on other mink farms.

*b) C2 variant*

The C2 variant was initially found in samples taken on 16^th^ Oct. 2020 from mink on farm 111 in southern Denmark (see Figure 2B) and was subsequently also found on farms 263 and 264 (samples taken on 11^th^ Nov. 2020) and on farm 274 (samples taken on 13^th^ Nov. 2020). These three farms were each located in central Jutland (Figure 2B). The mink sequences clustering within C2 belong to the pangolin lineage B.1.1.219, which has the lineage-defining changes encoding the substitution F157L in the S protein. All the sequences within C2 have this same change and, in addition, the viruses from farms 263, 264 and 274 have an additional change in the S protein coding sequence to produce the substitution A845S. The changes F157L and A845S have not been found in other Danish mink sequences. However, there are, in the GISAID EpiCoV database [21], 92 sequences from humans in DK belonging to the B.1.1.219 lineage in samples collected between 7^th^ Sept. 2020 and 1^st^ Feb. 2021.

*c) C3 variant*

The C3 variant was identified from three mink farms in central Jutland (Figure 2B). The first finding of this variant was in samples collected on 26^th^ Oct. 2020 from farm 166. Subsequently, the C3 variant was also found in samples from farm 156 taken on 23^rd^ Oct. 2020 and in samples from farm 186 taken on 29^th^ Oct. 2020. The C3 variant belongs to pangolin lineage B.1.1.170, which has no changes in the S protein compared to the index mink strain. In two of the sequences from farm 166, a change in the S protein gene resulted in G1167S. This change was not seen in the other two farms with a C3 variant and has not been found in other mink sequences within DK. There are in the GISAID EpiCoV database a total of 222 sequences from humans in DK belonging to the B.1.1.170 lineage in samples collected between 20^th^ Jul. 2020 and 8^th^ Feb. 2021.

*d) C4 variant*

In total, the C4 variant was found on seven mink farms in southern Denmark in the period from 16^th^ Oct. 2020 to 18^th^ Nov. 2020, and is the variant, from among the new virus introductions (lacking Y453F), that was seen on most farms during the mink outbreak. The C4 variant was first found in samples taken on farm 110 in southern Denmark. Subsequently, the C4 variant was also found on farms 223, 224, 225, 276, 283 and 288. The C4 variant belongs to pangolin lineage B.1.536, which does not contain specific lineage-defining changes in the S protein gene. However, this lineage has two lineage-defining changes in the sequence encoding the ORF3a protein (T229I) as well as in the N-protein gene (resulting in G215V), which were also found in 2 out of 3 sequences from farm 110. These changes were also seen in samples from other farms with the C4 variant. There are 42 sequences from humans registered in the GISAID EpiCoV database [21] from lineage B.1.536. They were all sampled from DK during the period 26^th^ Oct. 2020 to 22^nd^ Feb. 2021.

The sequences from farm 223 and farm 276 also have a change in the S protein in the receptor-binding domain at position N501T. This change in the S protein is present in 132 mink sequences in GISAID, of which three are from DK (from 4^th^ Nov. 2020), five are from Latvia (from April and July 2021), five from the NL (April to June 2020), five from Spain (from July 2020) and the remainder from the USA (from August and September 2020). The N501T variants from the other countries occur in different virus lineages. Like the Y453F change, the N501T substitution is at the interaction site between the S protein and the ACE-2 receptor [14]. This suggests that N501T is an adaptive change that occurred following infection of mink, in a similar way to the Y453F change from farm 1. The N501T change has also been found in 24 human sequences in DK that were sampled in the period 9^th^ Nov. 2020 to 29^th^ Mar. 2021 according to the GISAID EpiCoV database [21], these belong to three different lineages, namely A.28, B.1 and C.2.1.

*e) C6 variant*

The C6 variant was detected, for the first time, in samples taken on 23^rd^ Oct. 2020 from farm 151 in northern Jutland. Mink sequences from this farm belong to pangolin lineage B.1.1.294, which has no changes in the S protein compared to the index case. In the GISAID EpiCoV database [21], 143 sequences from humans in DK belonging to the lineage B.1.1.294 have been registered, the first of which was found in a sample from 14^th^ Sept. 2020. The C6 variant has not been found on other mink farms.

*f) C7 variant*

The mink sequences in C7 were found in samples collected on 12^th^ Nov. 2020 from farm 267 in southern Denmark. The C7 sequences belong to pangolin lineage B.1.1.159. This lineage also does not contain specific lineage-defining changes in the S protein sequence. A total of 40 human sequences from DK are registered in the GISAID EpiCoV database [21] for lineage B.1.1.159, of which the first sequence is from 21^st^ Sept. 2020 and the last is from 23^rd^ Nov. 2020. In one of the three mink sequences from farm 267, there is a change in the N-terminal region of the S protein (G21624C; R21T). This R21T change is not recorded in the human sequences of lineage B.1.1.159 in the GISAID EpiCoV database [21].

*g) C8 variant*

The C8 variant was first identified in mink from farm 217 in northern Jutland in samples taken on 2^nd^ Nov. 2020. This variant belongs to lineage B.1.177 that has only 4 lineage-defining changes in the entire SARS-CoV-2 genome, one of which results in the S protein change D614G (found in all the mink viruses in DK). In addition to this change, the sequence from farm 217 also encodes the substitution A222V in this protein. There are 1207 sequences from humans in DK within the B.1.177 lineage in the GISAID EpiCoV database [21] that were collected from 3^rd^ Aug. 2020 to 10^th^ May 2021.
