## Supplementary figures and images for "Emergence and spread of SARS-CoV-2 variants from farmed mink to humans and back during the epidemic in Denmark, June-November 2020"

### Supplementary Figure S1

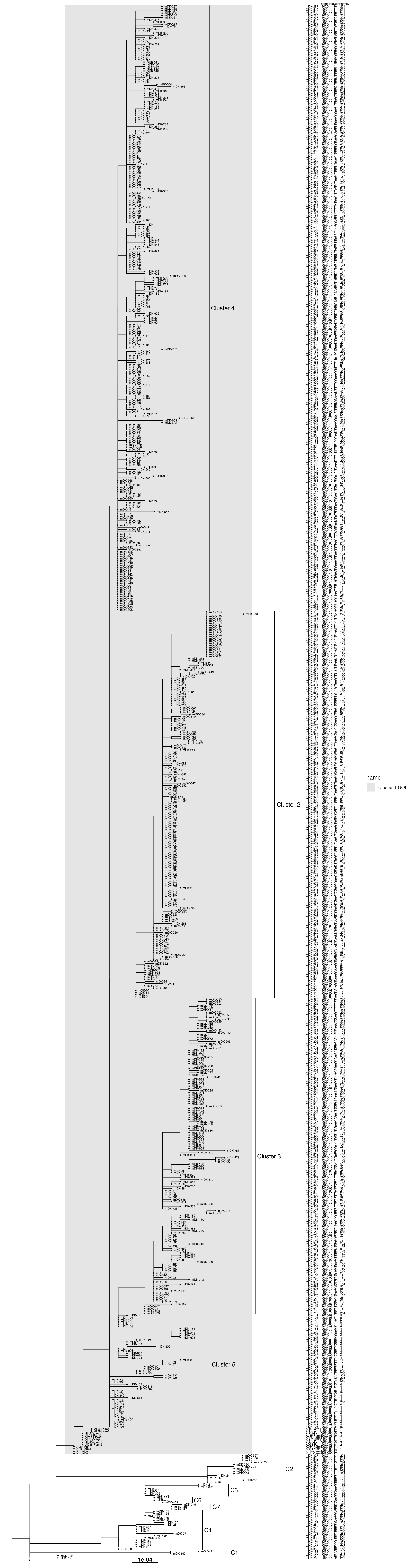
