## Supplementary Figure S2 for "Emergence and spread of SARS-CoV-2 variants from farmed mink to humans and back during the epidemic in Denmark, June-November 2020"

**Rasmussen et al. Mink Phylogeny Supplementary Figure S2**
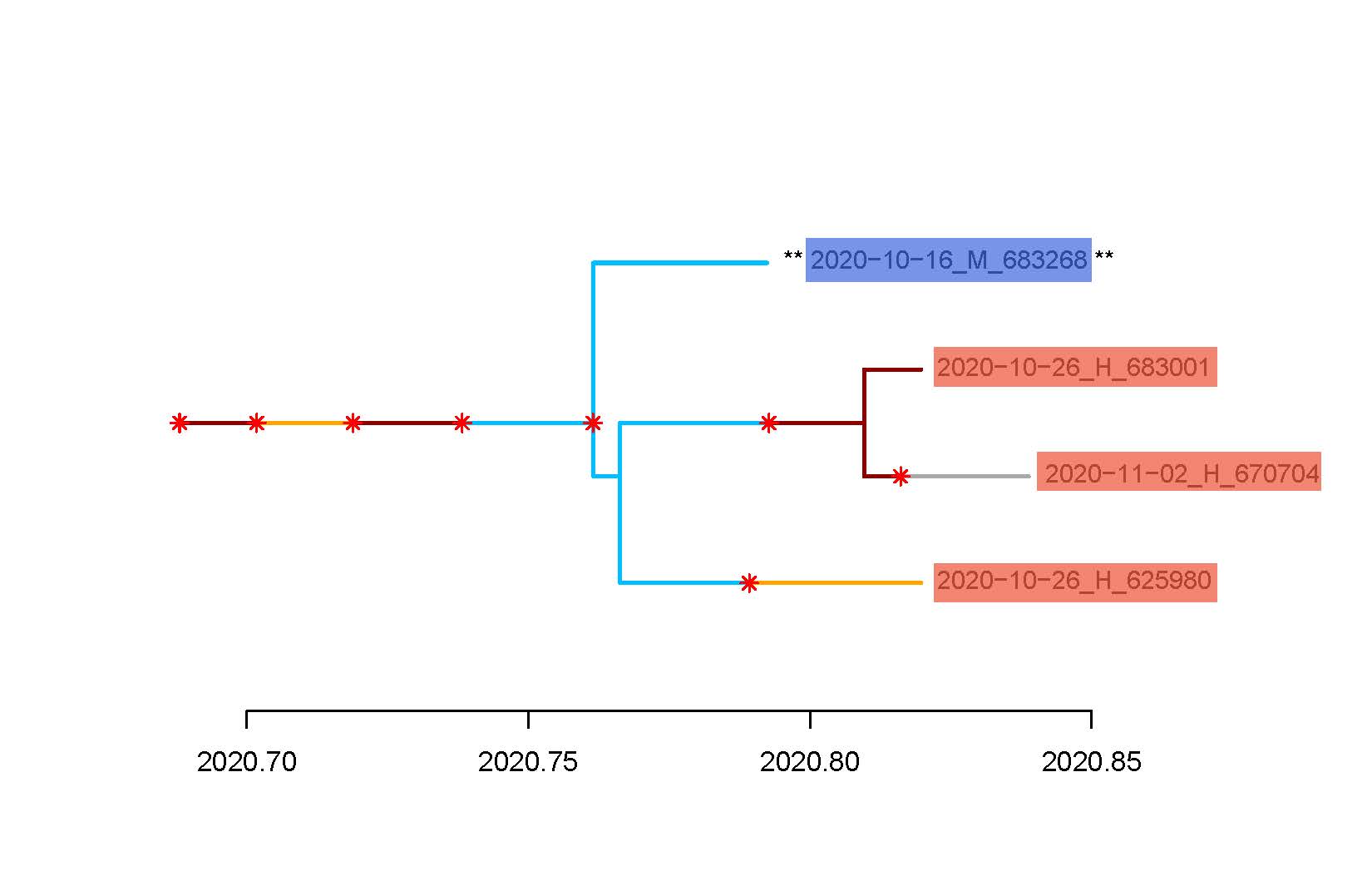

**Supplementary Figure S2.**  Colored phylogeny, showing inferred chain of SARS-CoV-2 transmission from a mink (sequence-name shaded in blue) to humans (names shaded in red). The phylogeny shown here is a detail of a larger inferred transmission tree inferred by the software TransPhylo [25]. Different branch colors correspond to different hosts, observed or un-observed, and a region of connected branches with a single color represents evolution happening inside a single host. Stars on branches correspond to transmission events. If a colored branch is connected to a leaf, then the host has been observed, otherwise it is an un-observed intermediate host inferred by TransPhylo. The mink host (branches colored blue, name shaded in blue) was sampled on October 10, and is inferred to have transmitted the virus to two humans, that were both sampled on October 26 (branches colored orange and brown). One of these human hosts was inferred to further have transmitted the virus to another human, sampled on November 2 (branch colored grey, sequence name: 2020-11-02_H_670704).
